## Supplementary Data for "Enhancing Portability of Trans-Ancestral Polygenic Risk Scores through Tissue-Specific Functional Genomic Data Integration"

**Supplementary Note 1**

We observed that the lead IMPACT transcription factor (TF) annotations had larger $\tau^*$ estimates than our lead TURF and TLand models for all traits tested. One explanation for this difference is the TURF and TLand models, on average, covered 4.6 times more common SNPs than IMPACT TF annotations. This increased coverage by TURF and TLand models leads to smaller per-SNP heritability estimates $(\hat{\tau})$ from stratified LDSC due to diluted heritability signal across a larger set of SNPs.

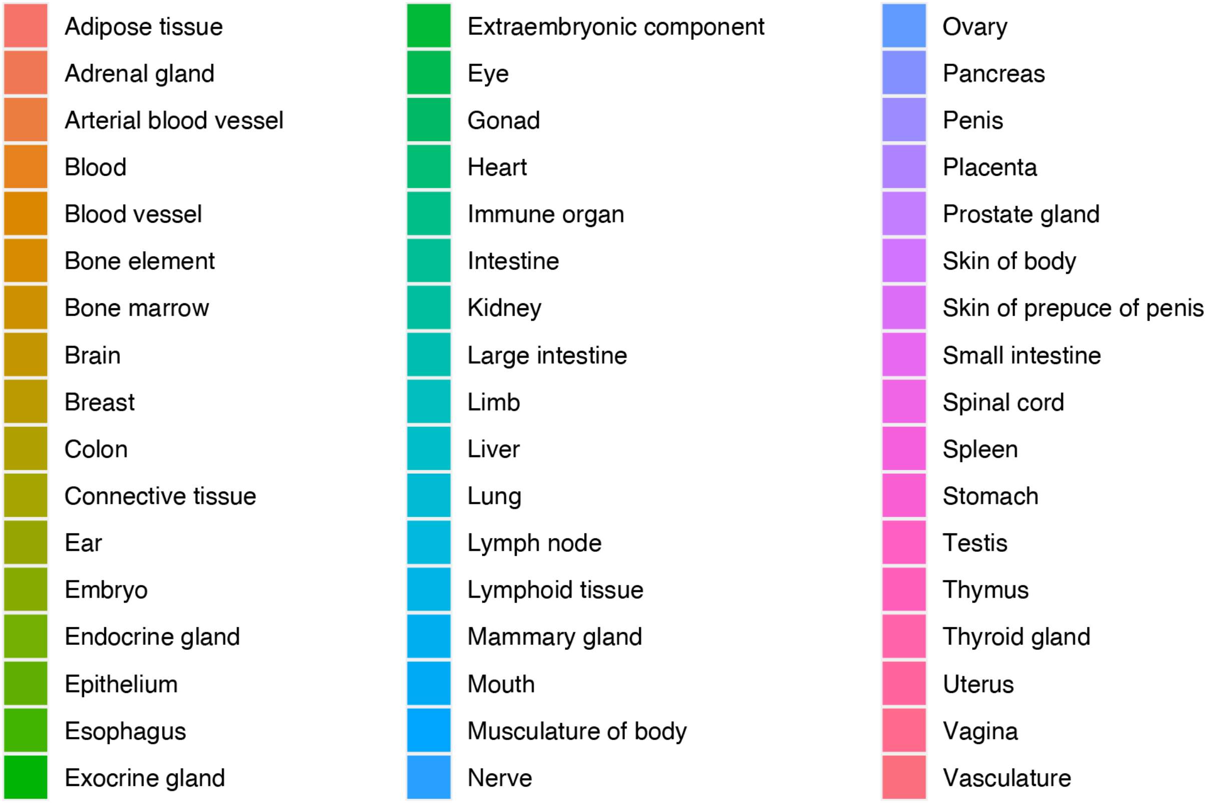


**Supplementary Figure 1 | Legend for tissue labels in TITR results - Figure 3 & Extended Data figures 1-2.**
