## Extended Data for "Enhancing Portability of Trans-Ancestral Polygenic Risk Scores through Tissue-Specific Functional Genomic Data Integration"

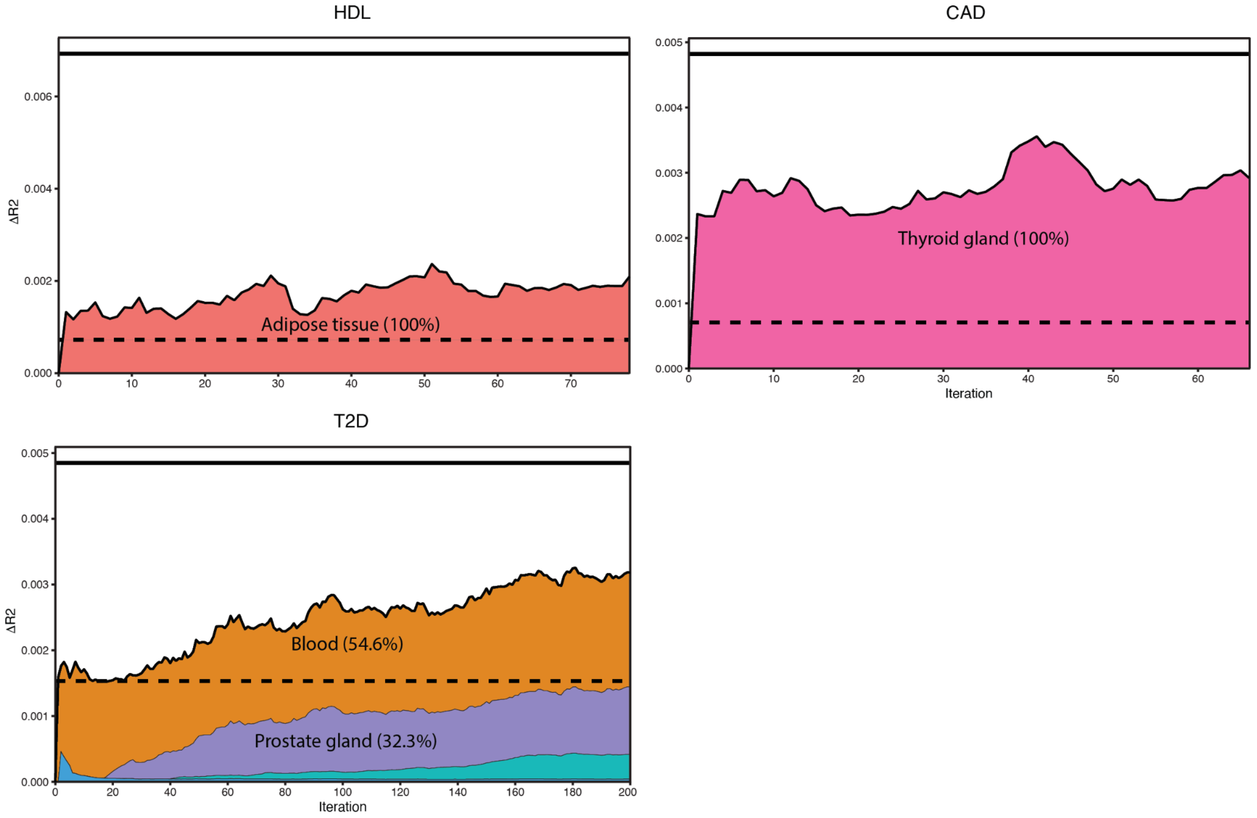


**Extended Data Figure 1 | Adjusted $R^2$ accuracy of EUR-to-AFR trans-ancestral PRS model for HDL cholesterol, coronary artery disease, and type 2 diabetes of the TITR optimization algorithm for TURF functional model.**

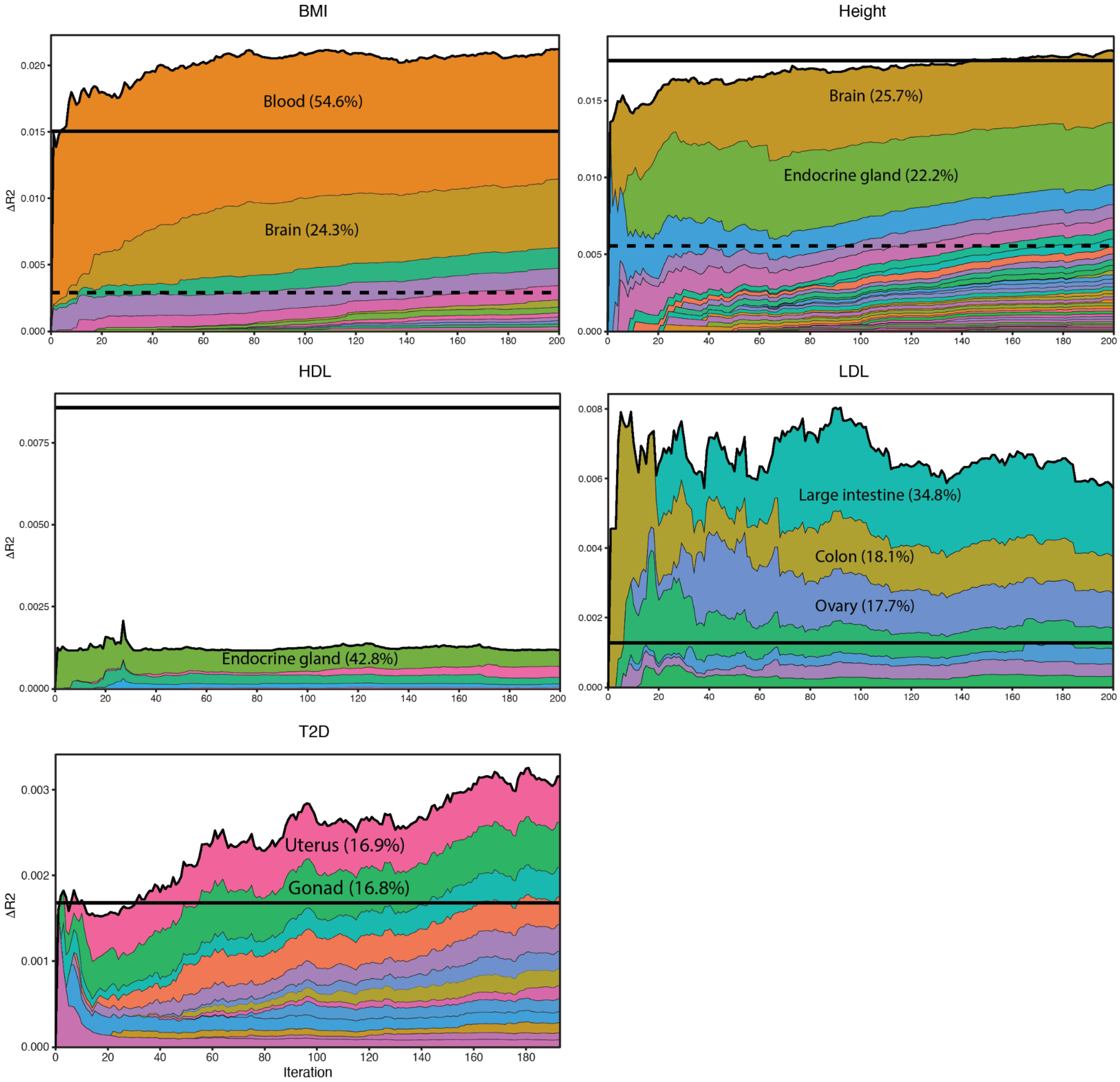


**Extended Data Figure 2 | Adjusted $R^2$ accuracy of EUR-to-AFR trans-ancestral PRS model for BMI, height, HDL cholesterol, LDL cholesterol, and type 2 diabetes of the TITR optimization algorithm for TLand functional model.**

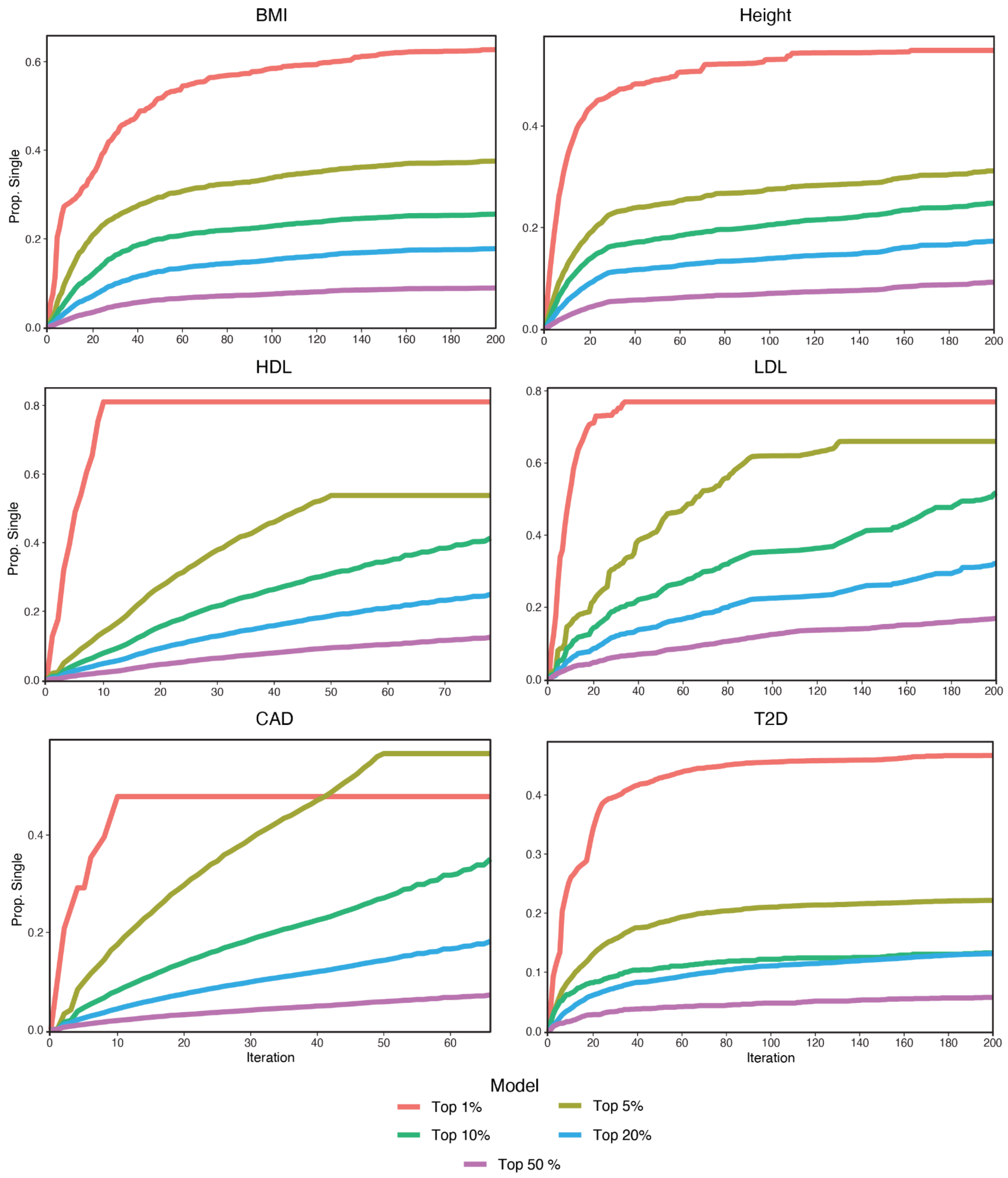


**Extended Data Figure 3 | Proportion of SNPs overlapping between TURF single tissue model (top 1/5/10/20/50% thresholds) and TITR PRS model for BMI, height, HDL cholesterol, LDL cholesterol, coronary artery disease, and type 2 diabetes.**

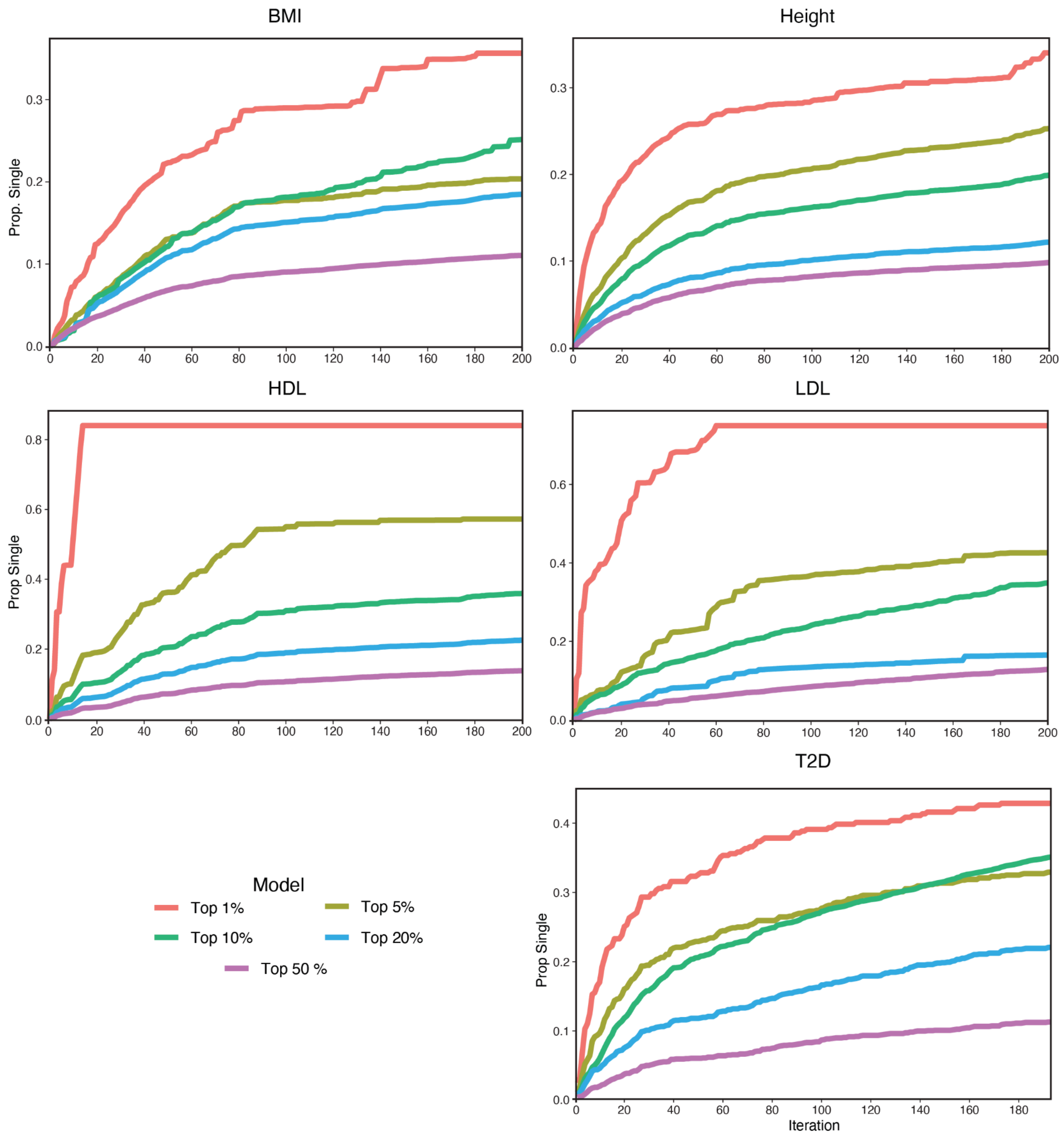


**Extended Data Figure 4 | Proportion of SNPs overlapping between TLand single tissue model (top 1/5/10/20/50% thresholds) and TITR PRS model for BMI, height, HDL cholesterol, LDL cholesterol, and type 2 diabetes.**
